# Plant coumarins enhance viral pathogenicity by miRNA-mediated apoptosis in insect herbivores

**DOI:** 10.1101/2025.09.29.679223

**Authors:** Jin-Yan Wang, Hao Zhang, Yuan Yuan, Mark A. Hanson, Wen-Li Xia, Jun-Xiang Zhou, Zhen-Dong Song, Jie-Xian Jiang, Xiang-Yun Ji, Nian-Feng Wan

## Abstract

Plant secondary metabolites (PSM) influence the susceptibility of insect herbivores to entomoviruses, but the apoptosis mechanism is not clear. Among plant hosts screened, the level of coumarin is correlated with NPV-mediated mortality, with high coumarin levels associated with high mortality and low coumarin levels with low mortality. Flow cytometry and morphological experiments showed that coumarins increased the apoptosis rate in NPV-infected larvae. miRNA transcriptomic analysis of coumarin-exposed larvae identified downregulation of miR-8 and miR-6497-3p, which negatively correlate with expression of pro-apoptotic kinases MAPK and ASK1. In addition, miR-9c-5p is upregulated, suppressing the apoptosis inhibitor 14-3-3 protein zeta. Our study suggests plant coumarins increase the susceptibility of larvae to NPV through miRNA-mediated regulation of apoptotic pathways. These findings suggest viral biocontrol efforts may be more successful on certain host plants, and provide molecular insight into how plant defense chemicals synergize with entomoviruses to control agricultural pests.

## INTRODUCTION

Plants generate secondary metabolites as a defence response against herbivore attack.^1^ Plant secondary metabolites (PSMs) (e.g., phenolics, nitrogen-containing organic compounds and terpenoids) have been shown to suppress the growth and reproduction of insect herbivores.^2,3^ Entomopathogens (e.g., entomoviruses) also commonly infect insect herbivores, and offer potential solutions for pesticide-free biocontrol.^4–6^ Up to now, the effects of PSMs on the susceptibility of insect herbivores to entomopathogens have not received great attention, although there have been wide reports on plant-mediated effects on such susceptibility.^7–9^ The molecular level explaining how plant metabolites interact with potential biocontrol agents, such as entomoviruses, is poorly understood.

Apoptosis serves as an important host defense mechanism for inhibiting virus replication by eliminating damaged cells.^10^ Among PSMs, plant coumarins are important lactone compounds distributed extensively in higher plants (e.g., Fabaceae, Rutaceae, Apiaceae, Asteraceae and Solanaceae). Coumarins exhibit high contact toxicity against lepidopteran larvae (e.g., cabbage caterpillars and diamondback moths),^11–13^ and cause host cell apoptosis.^14,15^ For example, plant coumarins induced the apopotosis of *Hyphantria cunea* larvae, alongside decreased body weight and lower expression levels of growth-related genes.^15^ Likewise, entomoviruses can also induce apopotisis in insect herbivores,^16^ such as the effects of nucleopolyhedroviruses (NPV) on *Spodoptera litura*^17^ and on *S. frugiperda*.^18^ Moreover, some entomoviruses have evolved mechanisms to subvert cellular apoptosis, facilitating virus replication, assembly, and release.^19,20^ However, the interaction of PSMs with entomovirus-induced apoptosis in insect herbivores has not been explored.

MicroRNAs (miRNAs), endogenous non-coding RNAs containing approximately 22 nucleotides.^21^ These short RNAs have emerged as important regulators of various physiological and pathological processes (e.g., apoptosis) by binding chiefly to the 3’-untranslated region (3’-UTR) of target mRNAs, where they exert their regulatory functions by mRNA degradation or translational inhibition.^22,23^ Previous studies have indicated that viral infections alter the expression of host miRNAs.^24^ For instance, Dengue virus (DENV) infection increased the expression of *Aedes albopictus* miR-252, which is proposed to target a DENV viral envelope protein, resulting in suppressed viral replication.^25^ A viral miRNA can further regulate lepidopteran host genes to promote viral transmission.^26^ In non-infectious contexts, miRNAs provide a mechanism to fine-tune host cell death and stress response pathways, including apoptosis via MAPK signalling,^27,28^ and host detoxification enzmyes.^29^ Insects also utilize the immune melanization response in viral control,^12,30^ which generates reactive oxygen species that can cause host cell damage and ensuing cell death and apoptosis.^31^ The regulation of these processes, which are modulated in concert upon viral infection, presents a challenge for the infected host: the host must manage the killing of infected cells, but must also prevent excessive autotoxic immune damage. To date, it remains unclear how host-targeting miRNAs might contribute to the response to PSM exposure, including the interaction of PSMs and entomovirus infection on host cell apoptosis.

The beet armyworm *Spodoptera exigua* (Lepidoptera: Noctuidae), a widely distributed polyphagous pest, is highly susceptible to baculoviruses (family Baculoviridae) such as SeMNPV (*Spodoptera exigua* multiple nucleopolyhedrovirus). In previous studies, we found that the mortality of SeMNPV-infected larvae differed among 14 plant species with highest mortality on *Glycine max*, intermediate on *Brassica oleracea*, and lowest on *Ipomoea aquatica*.^8^ By using metabolomic analyses of these three plant species, we found that four PSMs (genistein, kaempferol, quercitrin, and coumarin) were relatively high, each capable of enhancing the susceptibility of *S. exigua* to SeMNPV, with coumarins having the most potent effect.^12^ To elucidate how coumarins synergize with SeMNPV to increase viral pathogenicity, we investigated the interaction among plant coumarins, *S. exigua*, and SeMNPV using integrated approaches including flow cytometry, miRNA transcriptomics, and functional assays. We propose that plant coumarins enhance NPV pathogenicity by modulating miRNA-mediated apoptosis pathways in insect hemocytes (Figure 1). Our results reveal how plant chemical defenses interact with viral biocontrol agents, providing molecular insights to optimize pest control strategies by aligning host plant chemistry with entomovirus applications.

**Figure 1.**
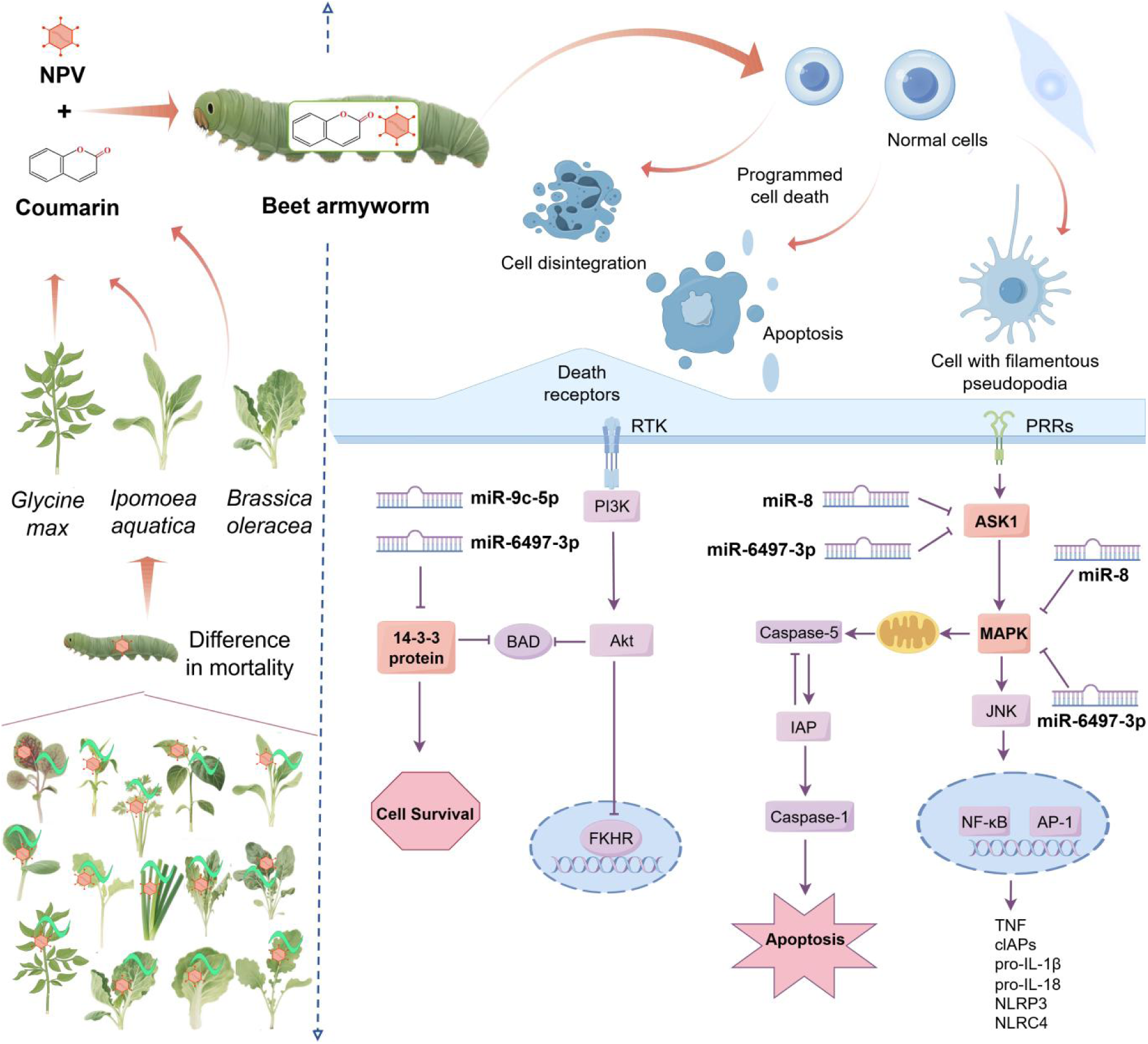
Proposed mechanism of plant coumarin-enhanced viral pathogenicity via miRNA-mediated apoptosis regulation.

## RESULTS

### Plants Secondary Metabolites Cause Differences in Susceptibility of *S. exigua* to SeMNPV

The susceptibility of *S. exigua* to SeMNPV (hereafter “NPV”) varied when larval *S. exigua* were fed on 14 different plant species, with the highest LD_50_ (lethal dose of 50% mortality, reported as log_10_-transformed dose per larva) on *I*. *aquatica* (2.97), intermediate LD_50_ on *B*. *oleracea* (2.63), and lowest LD_50_ on *G. max* (1.81) (Figure 2A). The results of metabolomic analyses indicated that the concentrations of four main plant phenolics (genistein, kaempferol, quercitrin, and coumarin) were higher in *G. max* than in *B. oleracea* or *I. aquatica* (Figure 2B). Among these four PSMs, coumarins had the greatest effect on enhancing the susceptibility of *S. exigua* to SeMNPV. Indeed, larval mortality 2-9 days post-treatment (dpt) was much higher in “coumarin+NPV” treatments (70.83%) compared to treatment with NPV (23.75%) or coumarin alone (47.5%) (Figure 2C).

**Figure 2.**
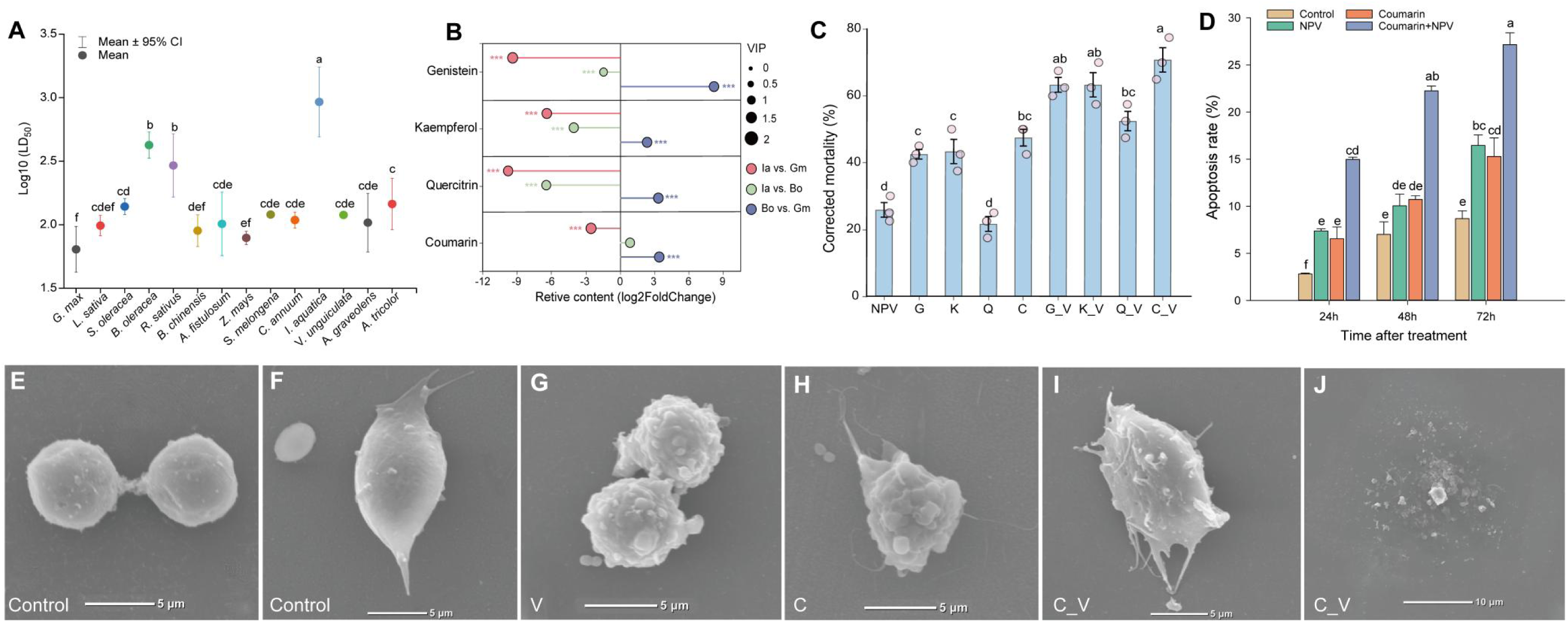
Plant-mediated susceptibility of caterpillars to entomoviruses and plant secondary metabolite-mediated differences in hemocyte apoptosis of entomovirus-infected caterpillars. (A) Mortality of beet armyworm larvae fed aliquots of SeMNPV on foliage of 14 crop plant species (data was available on zenodo: https://zenodo.org/records/16411366).^8^ Lower LD_50_ indicates greater mortality per dose. (B) Plant secondary metabolite (genistein, kaempferol, quercitrin, and coumarin) contents in three plants (data was available on zenodo: https://doi.org/10.5281/zenodo.14039264).^12^ Ia (*I. aquatica*), Gm (*G. max*), Bo (*B. oleracea*). Relative quantitative means of plant secondary metabolites were compared between each plant hots, with log-fold difference shown; for instance, a negative relative content of Ia vs Gm indicates Ia has less than Gm. *** indicates p < 0.001 (Student’s t-test) and OPLS-DA (VIP > 1). (C) Corrected mortality of *S. exigua* larvae treated with SeMNPV and plant secondary metabolites (data was available on zenodo: https://doi.org/10.5281/zenodo.14039264). V, NPV (2.12 OBs larva); G, genistein; K, kaempferol; Q, quercitrin; C, coumarin. G_V, genistein +NPV; K_V, kaempferol + NPV; Q_V, quercitrin+NPV; C_V, coumarin + NPV. Corrected mortality indicates increase in mortality relative to unchallenged controls. (D) Hemocyte apoptosis rate of caterpillars raised on different diets. (E-J) example hemocytes visualised with scanning electron microscopy inclduing control group granular oenocytoid (E) and plasmatocyte (F), hemocytes exposed to NPV alone (G), coumarin alone (H), or NPV + coumarin (I, J). Also see Table S5.

### Effects of Coumarin on the Apoptosis of NPV-infected *S. exigua* hemocytes

The apoptosis of larval *S. exigua* hemocytes (blood cells) was significantly impacted by the addition of coumarin to the diet for given NPV doses (no-NPV controls (0), and 2.12 × 10^8^ OB·mL^-1^) with a consistent trend in the effect of treatment over time (Figure 2D, Table S5). Combined treatments of NPV and coumarin resulted in significantly higher apoptosis rates compared to coumarin or NPV alone (*P*<0.001) (Figure 2D). Morphological observation of hemocytes by scanning electron microscopy confirmed that the control group hemocytes maintained a normal cellular structure (Figure 2E, F). However, exposure to coumarin, NPV, or both led to increasing disruption of hemocyte cellular structure, including the presence of apoptotic bodies, which were readily observed in a subset of cells on all diets 72 hours after NPV treatment (Figure 2G-J, also see Figure S3). We measured apoptosis rate by flow cytometry, and observed that the apoptotic bodies and filamentous pseudopodia in the coumarin+NPV treatment were much more common than those in the individual treatments fed either coumarin or NPV alone 72hpt (hours post treatment). In addition, cell disintegration was more readily observed in the coumarin+NPV 72hpt.

### Comparative miRNA sequencing of *S. exigua* treated with coumarins and NPV

In a previous study,^12^ we analyzed the differentially expressed genes (DEGs) of *S. exigua* after PSM and virus exposure. Here, we have utilized miRNA sequencing of *S. exigua* larvae to reveal if or how host miRNAs might respond and help to regulate host responses to coumarin exposure or viral infection. For control, 18.6 μg coumarin alone (C), 2.12 OBs·larva^-1^ of NPV alone (V), and coumarin+NPV combined treatment (C_V), we identified identified 3208,2180,1948 and 680 mature miRNAs on average. There were 45 DEGs (Differential Expression of miRNA Genes) in the control vs. V group, 34 DEGs in the V vs. C_V group, 95 DEGs in the control vs. V group, 2 DEGs in the V vs. C group, and 33 DEGs in the control vs. C_V group (|log_2_FoldChange| > 1, *P*-value < 0.05) (see Figure 3, Figure S3 and Table S2). Of these, 93, 45, and 2 DEGs were unique to the control vs. V, control vs. C, and V vs. C_V, comparisons respectively (Figure 3F).

**Figure 3.**
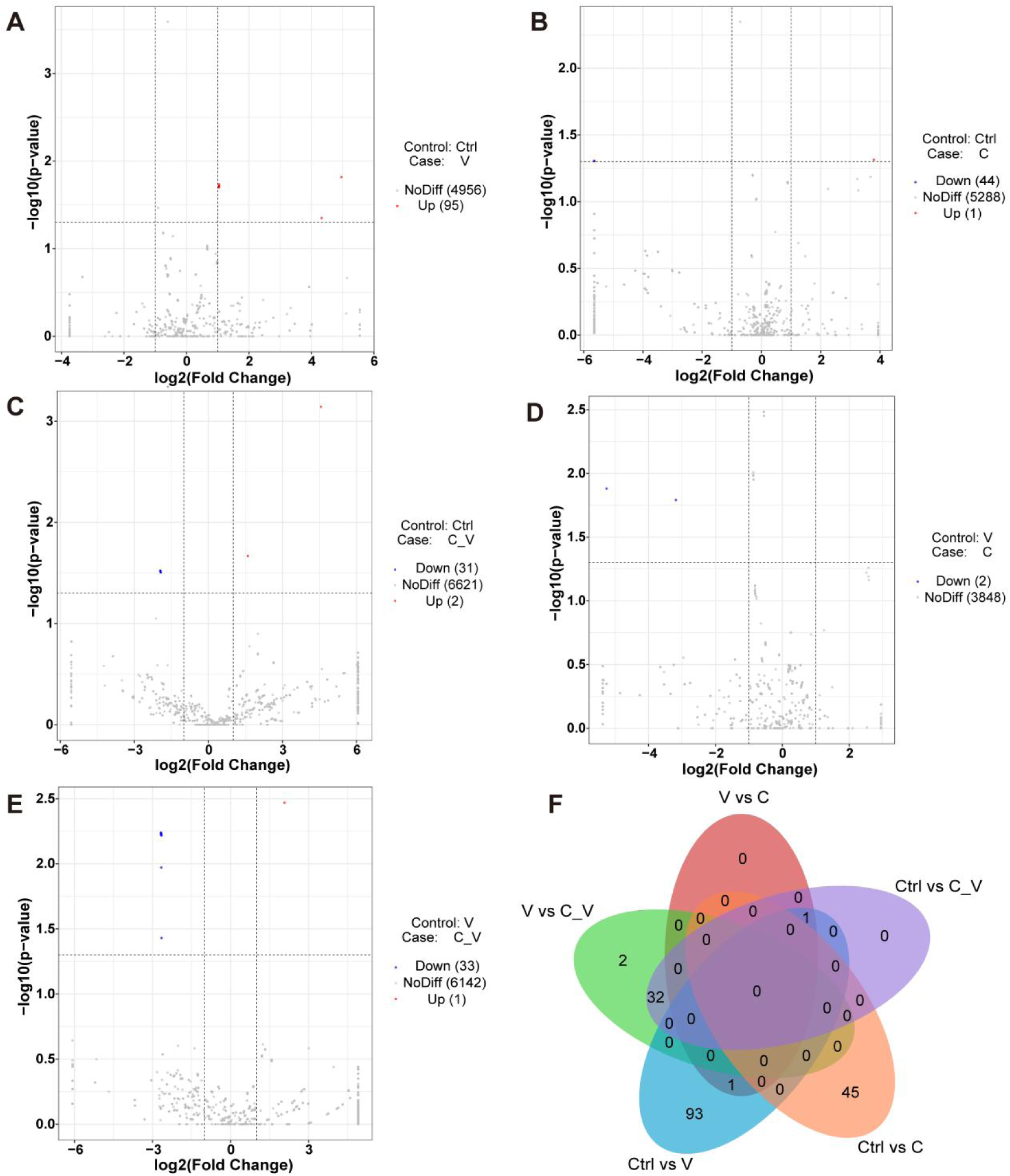
Differentially expressed miRNA gene (DEG) counts of *S. exigua* larvae treated with NPV and coumarin. Genes were defined as significantly differentially expressed if |log_2_Fold Change| > 1, and P-value < 0.05. (A-D) Volcano plots showing the up-regulated, down-regulated and unaffected genes between NPV treated and “NPV + coumarin” treated groups. (E) Venn diagram denoting DEGs that overlap or are unique among the four groups; Ctrl = Control; V = NPV; C = coumarin; C_V = coumarin+NPV. See Tables S1 and S2.

In the control vs. V, control vs. C, control vs. C_V, V vs. C, and V vs. C_V groups, there were 46, 14, 121, 28, 135 DEGs, with 27, 9, 33, 11, 31 DEGs impacting apoptosis-related pathways in gene ontology (GO) and kyoto encyclopedia of genes and genomes (KEGG) analyses (Figures 4A, B, S1 and S2). We used miRanda (v3.3a) to perform target gene prediction on the differentially expressed miRNA sequence. Interestingly, we identified three differentially expressed miRNAs that could theoretically regulate apoptosis directly. These included miR-8, miR-6497-3p, and miR-9c-5p that are predicted to target Mitogen-Activated Protein Kinase (MAPK), miR-8 and miR-9c-5p are also predicted to target Apoptosis Signal-Regulating Kinase (ASK1), and miR-6497-3p and miR-9c-5p are also predicted to target to 14-3-3 protein zeta (Figures 4C and S3). Accordingly, MAPK, ASK1, and 14-3-3 protein zeta were all differentially expressed in control vs. coumarin, NPV, or coumarin+NPV treatments (Figure 5).

**Figure 4.**
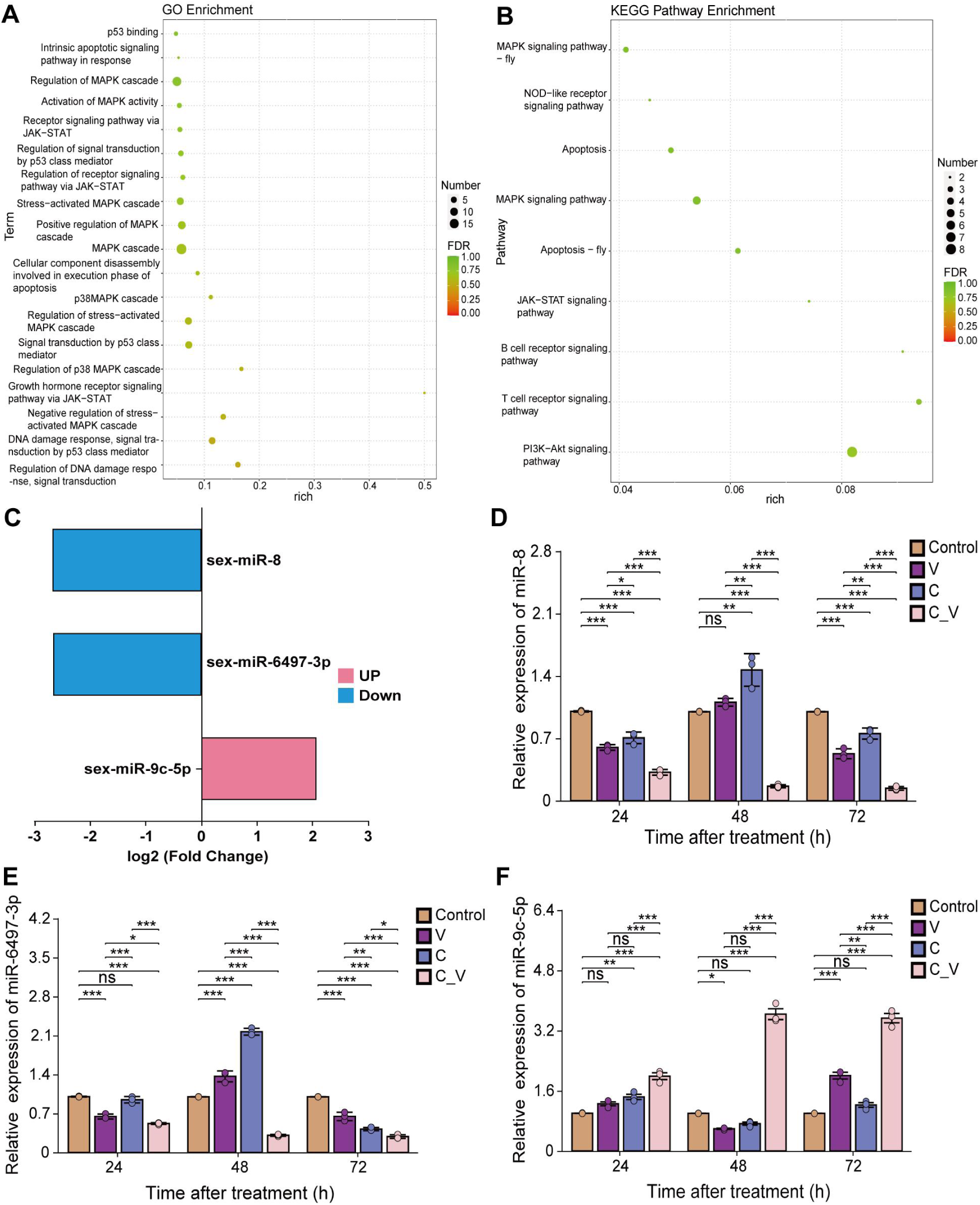
GO and KEGG pathway enrichment on the target apoptosis-related genes of differentially expressed miRNA and miRNA expression of *S. exigua* larvae. V = NPV; C = coumarin; C_V = coumarin+NPV. Enrichment factor refers to the ratio of the number of differentially enriched genes relative to the number of annotated genes in a pathway, with a larger enrichment factor indicating greater enrichment. False disocvery rate (FDR) generally ranges from 0 to 1, with FDR closer to zero indicating more significant enrichment. (A) GO enrichment on the target apoptosis-related genes of differentially expressed miRNA of *S. exigua* larvae in V vs. C_V groups. (B) KEGG pathway enrichment on the target apoptosis-related genes of differentially expressed miRNA of *S. exigua* larvae in V vs. C_V groups. (C) miRNA expression of *S. exigua* larvae in V vs. C_V group. (D-F) Relative expression of miRNA in four groups (control, NPV, C, C_V). (D-F) relative expression of indicated miRNAs given different treatments at 24, 48, and 72hpt. Also see Figures S1-S3, S5 and Tables S3.

**Figure 5.**
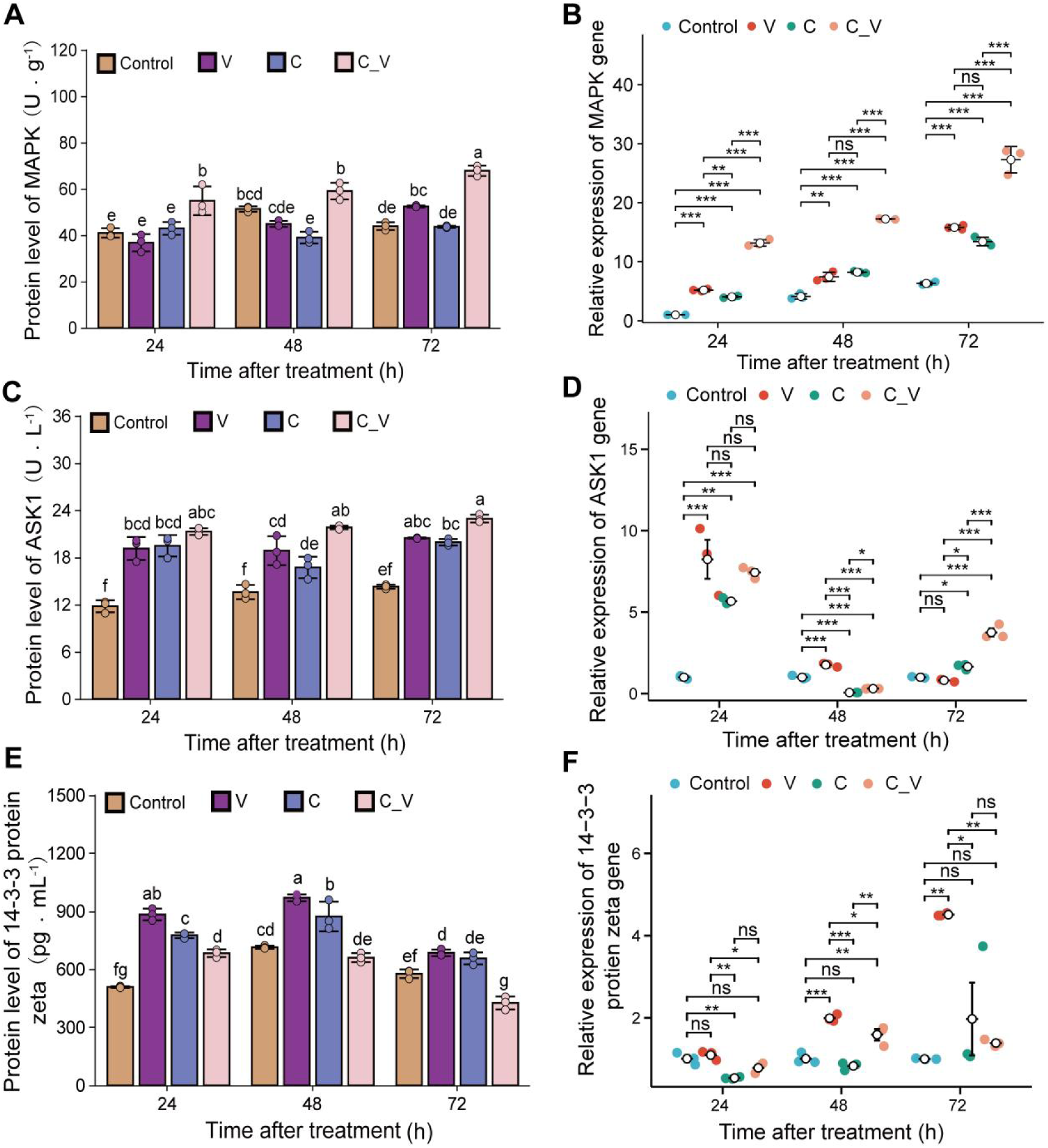
The enzyme protein levels and relative gene expression levels of *S. exigua* in different groups. (A, C, E) Enzyme protein levels of MAPK, ASK1 and 14-3-3 protein quantified by ELISA. (B, D, F) The relative gene expression levels of MAPK, ASK1 and 14-3-3 protein by qRT-PCR. V = NPV; C = coumarin; C_V = coumarin+NPV. See Tables S3 and S5.

### Effects of coumarins on the gene expressions and enzyme activities of NPV-infected *S. exigua*

Given the signature of DEGs relating to apoptosis, alongside differential expression of three miRNAs predicted to target regulators of apoptosis, and differential expression of the genes these miRNAs would putatively target, we chose to characterize these three miRNAs further. Gene expression levels of miR-8, miR-6497-3p, and miR-9c-5p were all affected by NPV viral infection, coumarin treatment, and the combination of coumarin+NPV. Indeed, in the coumarin+NPV treatment, levels of miR-8 and miR-6497-3p were consistently reduced, and levels of miR-9c-5p were consistently increased (Figure 4C-F). Furthermore, good concordance was seen for the enzyme activities (Figure 5A, C, and E), and expression levels of the target genes (Figure 5B, D and F) with effects that were consistent in direction for each treatment relative to the miRNAs proposed to regulate them (also see Table S5). In coumarin+NPV treatments, MAPK and ASK1 protein and mRNA levels were consistently higher than control, or coumarin or NPV alone, and 14-3-3 protein zeta levels were consistently depressed relative to coumarin or NPV treatments, indicating a failure of this gene to be induced upon coumarin+NPV treatment (Figure 5).

### Validation of the regulation between miRNAs and their target genes

We observed a good correlation between the expression of these miRNAs and their putative target genes implicated in apoptosis. We next used a dual-luciferase assay to verify binding of these miRNAs to their putative target sequences. Cotransfection of the MAPK-wild-type plasmid (MAPK-wt) with miR-8 or miR-6497-3p mimics led to a significant decrease in luciferase activity (*P*<0.001, Figures 6A and D), indicating successful binding of the miRNAs to the MAPK-wt sequence. In contrast, cotransfection of the MAPK-mutant plasmid (MAPK-mu) with miR-8 or miR-6497-3p mimics did not result in a significant change in luciferase activity, indicating a specific activity for only wild-type MAPK sequence. The same pattern was observed for ASK1 with miR-8 or miR-9c-5p, and for the 14-3-3 protein zeta with miR-9c-5p or miR-6497-3p (Figures 6B, C, E and F). We next injected *S. exigua* larvae with synthetic agonist miRNAs (AgomiR) to mimic an overexpression-type scenario. Independent RT-qPCR results showed that injection of miR-8 or miR-6497-3p significantly reduced the relative expression of the MAPK gene, injection of miR-8 or miR-9c-5p decreased the relative expression of the ASK1 gene, and injection of miR-6497-3p or miR-9c-5p reduced the relative expression of the 14-3-3 protein zeta gene. Conversely, injection of antagonistic miRNAs that should inhibit the target miRNAs significantly increased the expression of the target genes MAPK, ASK1, and 14-3-3 protein zeta (*P*<0.001, Figures 6G-I).

**Figure 6.**
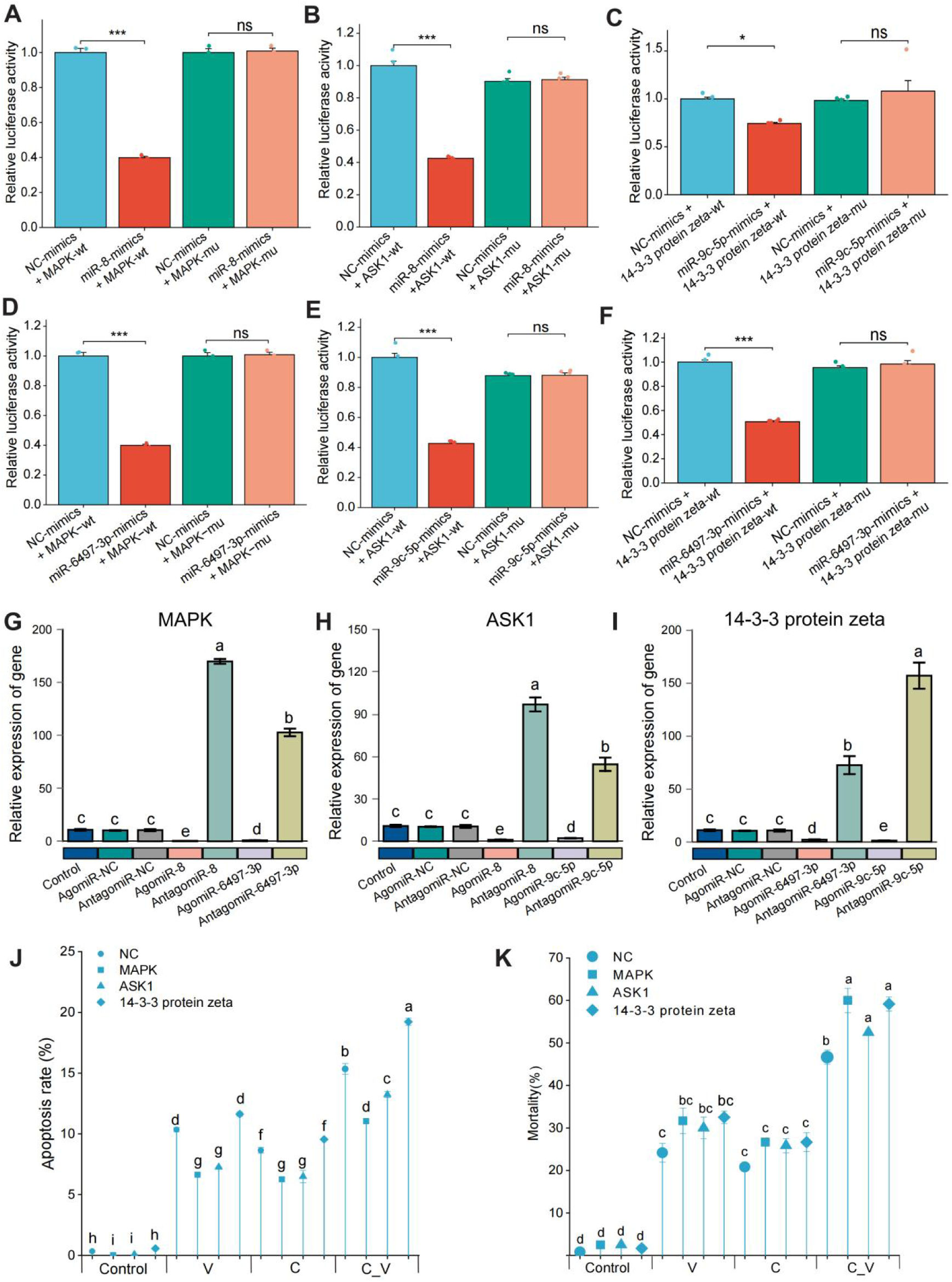
Changes in fluorescence activity of double luciferase reporters and relative gene expression, apoptosis, and mortality of *S. exigua* larvae treated with the synthesized miRNA and siRNA. (A-F) fluorescence activity of double luciferase reporters for target miRNAs. –wt and –mu indicates wild-type and mutant docking sites. (G-I) The relative gene expression levels of MAPK, ASK1 and 14-3-3 protein zeta of *S. exigua* larvae treated with the synthesized miRNA. (J) The apoptosis of *S. exigua* larval hemocytes treated with the synthesized siRNA. (J) The mortality of *S. exigua* larvae injected with the synthesized siRNA of genes (NC control, MAPK, ASK1 and 14-3-3). V = NPV; C = coumarin; C_V = coumarin+NPV. See Figure S4, S5 and Tables S1-S4.

These results indicate that these miRNAs affect the expression of MAPK, ASK1, and 14-3-3 protein zeta expression in important ways. However, the logic of target gene expression changes following agonist or antagonist miRNA injection was not always consistent with expectations. It is likely that the millieu of induced miRNAs and regulatory changes taking place during coumarin exposure and/or viral infection has important impacts on fine-tuning the expression of these target genes, which were not balanced in the same way in our experiments requiring artificial injury to inject miRNAs. Nevertheless, these findings collectively suggest that the three miRNAs (miR-8, miR-6497-3p, and miR-9c-5p) have the potential to regulate expression levels of MAPK, ASK1, and 14-3-3 protein zeta in vivo.

### Functional analyses of proteins in apoptosis and mortality

So far we have demonstrated that miRNAs miR-8, miR-9c5p, and miR-6497-3p regulate the expression of target genes MAPK, ASK1, and 14-3-3 protein zeta. To investigate whether these target genes have an importance in the plant-dependent susceptibility of *S. exigua* to NPV, we used RNA interference (RNAi) to knock down the expression of these three candidate genes and monitored hemocyte apoptosis and larval mortality. RNAi gene silencing reduced the expression levels of the target genes by 60% to 74% in the siRNA-treated larvae at 12 and 24 hours post-treatment, compared to the control (Figure S5A). Inhibition of MAPK and ASK1 significantly suppressed the apoptosis rate compared to that of controls (*P<*0.005 in all cases). Conversely, the apoptosis rate increased following the inhibition of 14-3-3 protein zeta expression in the coumarin+NPV group (*P<*0.0001) (Figure 6J and K). Furthermore, larval mortality of *S. exigua* treated with coumarin and NPV alone, or in combination, was significantly elevated after MAPK, ASK1, and 14-3-3 protein zeta genes were silenced by RNAi (*P*<0.0001). Gene knockdown had no significant effect on the larval mortality in the control, coumarin, or NPV groups (*P*=0.285 to 1.000) (Figure 6K).

In conclusion, miRNAs miR-8, miR-9c5p, and miR-6497-3p regulate the expression of MAPK, ASK1, and 14-3-3 protein zeta. In turn, MAPK, ASK1, and 14-3-3 protein zeta levels are key determinants of larval mortality and apoptosis following coumarin or NPV exposure. Moreover, the combination of coumarin and NPV is greater than either effect individually, indicating a non-redundant impact on host mortality. Taken together, the PSM coumarin enhances the NPV-mediated mortality of *S. exigua,* highlighting the potential importance of considering PSMs, and accordingly host plant, in the effective application of entomoviruses to pest insects.

## DISCUSSION

Differences in the susceptibility of insect herbivores to entomoviruses can be attributed to multiple factors, including the growth and behavioral responses of insect herbivores to different host plants,^8,32,33^ the immune responses of herbivores to entomoviruses,^34–36^ plant-mediated alterations of insect gut physiology (particularly the peritrophic membranes), the impact of plant trichome architecture on entomovirus infection,^37–39^ and the diversity of phytochemical profiles across various plant species.^40,41^ Our recent studies indicate that PSMs also affected such susceptibility, and plant coumarins help to increase the susceptibility of insect herbivores to entomoviruses.

Host plants with high content of plant coumarins are correlated with increased susceptibility of *S. exigua* to NPV.^12^ In this study, we have furthered the understanding of this interaction by detailing the impact of coumarin on apoptosis regulatory mechanisms. We found that coumarin alone induced miRNAs regulating host cell apoptotic pathways, and these same miRNAs are induced by NPV viral infection. Apoptosis often plays a “double-edged sword” role in virus-host interactions, as higher apoptosis rates can prevent viral proliferation if host cells are destroyed before viruses can effectively replicate, but this incurs damage to the host, and can ultimately promote viral infection if the virus is not cleared at an early stage.^42–44^ Thus, the pro-apoptotic effect of coumarin may contribute to overwhelming the host’s ability to effectively clear infecting virus, increasing the chance the host is cut down by its own double-edged sword, yielding additive or synergistic mortality outcomes.

Bideshi et al.^45^ reported that a virus in *S. frugiperda* encodes a caspase that triggers apoptosis in Sf21 cells, thereby promoting viral replication and spread. Similarly, Xu et al.^46^ found that inhibiting pro-apoptotic factor (R1PK1) reduces pro-inflammatory factors (R1PK3 and MLKL) and lowers the viral load in the host. Interestingly, recent studies have suggested an involvement of the insect melanization immune response in controlling viral infection.^12,30^ The melanization response involves the action of prophenoloxidases by macrophages and granular oenocytoids, which in turn generates reactive oxygen species, some of which can help to direct tissue repair programs, or participate in destruction of invading organisms.^47^ It would be interesting to confirm how PSMs such as coumarins, or viruses such as NPV, might impact the regulation of the insect melanization response, possibly affecting the activation of apoptotic pathways.

miRNAs are involved in coordinating a variety of cellular processes by regulating downstream genes, such as cell proliferation,^48^ morphogenesis,^49^ and metabolism.^50^ As miRNAs could participate in the process of normal cell physiology, including cell differentiation, apoptosis, proliferation, and cell cycle arrest,^51,52^ identifying apoptosis-related miRNAs during viral infection is essential to explore the fine-tuning of apoptosis mechanisms that prevent excessive host self-damage. In this study, we identified miRNAs consistently differentially expressed across different coumarin or virus infection treatments. We found that the target mRNAs of these differentially expressed miRNAs were, to some extent, enriched for apoptosis pathway gene ontologies. We identified and validated the effects of three miRNAs (sex-miR-8, sex-miR-9c-5p, and sex-miR-6497-3p) that impact the expression of apoptosis-regulating genes. miR-8 is a member of the miR-200 family of miRNAs, known to regulate cytoskeletal organization, epithelial cell proliferation, and apoptosis.^53–55^ The family of miR-9-5p has further been implicated in juvenile hormone biosynthesis and response to polyphenols in locusts and carp, respectively.^56,57^ Although the role of miR-6497 in cell apoptosis has not received wide attention, Liu et al.^58^ revealed that bmo-miR-6497-3p regulates circadian clock genes in silkworms.

In this study, the targeting of Mitogen-Activated Protein Kinase (MAPK), Apoptosis Signal-Regulating Kinase (ASK1), and 14-3-3 protein zeta, underscores the role of miRNAs in the apoptosis signaling pathway. Our dual-luciferase assays demonstrated that both miR-8 and miR-6497-3p inhibited MAPK expression, both miR-8 and miR-9c-5p inhibited ASK1, while both miR-6497-3p and miR-9c-5p inhibited 14-3-3 protein zeta. This molecular interaction is crucial, as MAPK and ASK1 are key signaling molecules participating in cell apoptosis,^59^ and 14-3-3 proteins are the inhibitors of apoptosis.^60,61^ We observed a decrease in miR-8, and miR-6497-3p upon coumarin and/or NPV treatments, but an increase in miR-9c-5p. Accordingly, we also saw an increased MAPK and ASK1 activity after coumarin and/or NPV treatments, but a depressed activity of 14-3-3 protein zeta after coumarin+NPV treatment. There is thus a consistency in the direction of these miRNAs being decreased and the mRNAs they regulate being increased, or vice versa, in vivo. It is therefore puzzling that agonist and antagonist miRNAs had a consistent direction of effect during miRNA injection experiments simulating overexpression. In this regard, we cannot exclude the possibility of secondary effects of miRNA manipulation, such as crude dysregulation of apoptosis by injection of miRNAs into the body cavity leading to a generic greater apoptotic rate. Alternatively, Zhang et al.^62^ reported that *Cnaphalocrocis medinalis* granulovirus regulates apoptosis by targeting AIF1 and ASPP1 through two miRNAs (tca-miR-3885-5p and tca-miR-3897-3p) to promote infection. Thus, it is likely that complex interactions of multiple miRNAs contributes to the in vivo signals seen in our gene expression and protein activity data. Nevertheless, our study shows the potent impact of the miRNAs miR-8, miR-6497-3p, and miR-9c-5p on the regulation of these apoptosis-regulating genes.

The effects of host plants or viral infections is often conducted with a focus on a specific compound or infectious agent. By consistently measuring individual and combined effects of coumarin and NPV treatments across our full set of experimental data, our study highlights the unique interactions that take place only during co-occurring treatment but not individual treatments. For instance, 14-3-3 protein zeta activity was induced in the coumarin+NPV treatment 24hpt, but ultimately was depressed relative to control larvae by 72hpt (Figure 5E). However, individual coumarin and NPV treatments retained a higher protein activity signal. Alongside the many observations of increased apoptosis rate and larval mortality, this result highlights the unique interplay that PSMs and entomoviruses can have on their insect hosts. As entomoviruses, including NPV, are promising biocontrol agents for targeting specific insect pests, our results highlight the utility of considering host plants and their PSMs in the application of NPVs; for instance, NPV may be most effective in controlling *S. exigua* when deployed on *G. max* or other host plants high in PSMs.

Our findings suggest a complex interplay between plant coumarins and molecular targets involved in the apoptotic response of larval *S. exigua* to NPV. The role of plant coumarin secondary metabolites extends beyond individual toxicity on insect herbivores, increasing their susceptibility to co-occurring viral infections. Here we found that plant coumarins are key modulators of specific miRNA expressions, particularly affecting miR-8, miR-9c-5p, and miR-6497-3p. Importantly, these microRNAs regulate apoptosis-related proteins, such as MAPK and ASK1, which are also important in the response to virus infection. Thus, our findings highlight how plant coumarins, defense chemicals, enhance the insecticidal efficiency of entomoviruses against agricultural pests through combinatory effects on host cell apoptosis. This study not only advances our understanding of plant-mediated effects on the interactons of insect herbivores and their entomopathogens, but also provides an unappreciated synergy of plant secondary metabolites and entomopathogens in controlling insect pests.

### Limitations of the study

A limitation of this study is that it focused on the PSM coumarin, whereas other PSMs can also contribute to plant-mediated synergistic effects on NPV mortality. In addition, the technical limitations of the model system required an invasive method of introducing miRNAs to the body cavity. In the future, gene editing approaches could allow endogenous genetic manipulations in *S. exigua* or other lepidopteran species, which can bypass the invasive injury incurred during injection of miRNAs or siRNAs. While flow cytometry, morphological techniques, and gene and protein level investigations provide valuable insights, the driving effects behind miRNA induction and their cause or effect relationship on apoptosis should be further explored.

## RESOURCE AVAILABILITY

### Lead contact

Further information and requests for resources and reagents should be directed to and will be fulfilled by the lead contact, Prof. (Dr.) Nian-Feng Wan.

### Materials availability

The plants, plant secondary metabolites, insect and SeMNPV lines used in this study will be made available upon request.

### Data and code availability

● The datasets of transcriptomes used in this study are available in the NCBI SRA database with the submission ID SUB15481248. Data on the mortality of beet armyworm larvae fed aliquots of SeMNPV on foliage of 14 crop plant species was available on zenodo: https://zenodo.org/records/16411366. Data on plant secondary metabolite (genistein, kaempferol, quercitrin, and coumarin) contents in three plants was available on zenodo: https://doi.org/10.5281/zenodo.14039264. Data on corrected mortality of S. exigua larvae treated with SeMNPV and plant secondary metabolites was available on zenodo: https://doi.org/10.5281/zenodo.14039264. Other data was deposited on zenodo: https://zenodo.org/records/16411394.
● This paper does not report original code.
● Any additional information, if required, is available from the corresponding author on reasonable request.

## ACKNOWLEDGMENTS

We thank the drawing tools provided by Figdraw. This study was sponsored by National Natural Science Foundation of China [grant number: 32502481], SAAS Program for Excellent Research Team [grant numbers: 2022[017]], and the National Ten Thousand Plan-Young Top Talents of China. For the purpose of open access, the authors have applied a ‘Creative Commons Attribution (CC BY) licence to any Author Accepted Manuscript version arising from this submission.

## AUTHOR CONTRIBUTIONS

N.F.W. and J.X.Y. conceived and designed the study; J.Y.W., M.A.H., J.X.Z., J.X.J. and N.F.W. wrote the manuscript; J.Y.W., H.Z., W.L.X., Y.Y., Z.D.S. and X.Y.J. conducted the experiments; J.Y.W., H.Z., J.X.J. and N.F.W. analyzed the data.

## STAR★METHODS

### EXPERIMENTAL MODEL AND SUBJECT DETAILS

#### Insects

*Spodoptera exigua* eggs were originally acquired from cabbage in the Fengxian District, Shanghai, China, in late September 2023. After undergoing surface sterilization with 5% formaldehyde, they were kept in clean chambers. The emerging larvae were fed on an artificial diet in climate chambers set at 27.0 ± 1.0 °C, 80.0 ± 5.0% relative humidity, and a 14:10-hour light-to-dark cycle. The artificial diet consisted of wheat germ, yeast powder, soybean powder, casein, agar, potassium sorbate, and L-ascorbic acid. Adults of *S. exigua* were provided with a 10% honey solution and raised in cylindrical plastic containers with paper for egg-laying. *S. exigua* was used in this study after being maintained for seven generations in the laboratory.

#### Coumarin and SeMNPV

Coumarin, sourced from Shanghai Yuanye Bio-Technology Co., Ltd., was incorporated into an artificial diet at a concentration of 0.1% of the diet’s dry weight. Larval S. exigua infected with NPV were homogenized in a glass homogenizer containing phosphate buffer solution (PBS) at 0.01 M and pH 7.0. The homogenate was then filtered through three layers of gauze, a process repeated four times to eliminate impurities and concentrate the virus. The filtrates underwent centrifugation twice at 900 × g for 15 minutes and twice more at 10,000 × g for 30 minutes to further purify the occlusion bodies (OBs). Subsequently, the purified OBs were subjected to density gradient ultracentrifugation and stored at 4 °C or subsequent use. During the experiment, the dosage of OBs was determined using a hemocytometer under a microscope at 400× magnification after the OBs had been diluted 100-fold. Ultimately, the NPV dosage was adjusted to 2.12 × 10^8^ OB·mL^-1^ using sterile water.

#### Coumarin and SeMNPV exposure

Early 3rd-instar larvae, (within 6 hours post-molt, were selected and placed into 48-well plates, with one larva per well, and starved for 6 hours. Each larva was then fed a 0.125 cm^3^ artificial diet cube treated with coumarin and SeMNPV in a factorial design. For the coumarin treatments, the artificial diet cube was either a PSM-free control or contained a dosage of 0.1% coumarin (186 μg per larva). For the NPV treatments, 1 μL of an NPV suspension (2.12 × 10^3^ OB·mL^-1^ (equivalent to 2.12 OBs per larva) or sterilized water for NPV-free controls was dispensed onto the artificial diet cube. After adding the cube, the plates were covered to prevent larval escape. To avoid NPV cross-infection, NPV-infected and uninfected larvae were isolated in climatic chambers in different rooms. Only larvae with similar body sizes that entirely consumed their diet cube within 24 hours, thereby taking a 100% dose of NPV (2.12 OBs per larva), were used in experiments.

#### The LD_50_ of SeMNPV to larvae of *S. exigua* feeding on 14 plant species

To evaluate the susceptibility of larval *S. exigua* to SeMNPV on different host plants, the LD_50_ (median lethal dose) was determined on the 14 plant species. A leaf disc bioassay was employed, where 1 µl of viral suspension was applied to 5 mm diameter leaf discs and allowed to dry. Each leaf disc was placed on a moistened filter paper disc (1 cm diameter) within the wells of a plastic bioassay plate. Three replicates were prepared for each virus concentration (14, 70, 350, 1750, and 8750 occlusion bodies per larva), with each bioassay repeated three times. Larvae that consumed the entire leaf disc within 24 hours (representing 100% of the applied viral dose) were individually transferred to sterile 18 ml plastic tubes containing fresh artificial diet and maintained at 27°C until death or pupation. Larval mortality was recorded daily, and LD_50_ values were calculated by fitting the data to an appropriate equation. Data from Wan et al. (2016).

#### Differential metabolites among host plants utilized by *S. exigua*

To quantify the levels of plant secondary metabolites (PSMs) in the target plants, ultra-high-performance liquid chromatography-quadrupole time-of-flight mass spectrometry (UHPLC-QTOF-MS) was used to analyze 18 leaf samples (2 g each) collected from cabbages (*Brassica oleracea*), soybeans (*Glycine max*), and water spinach (*Ipomoea aquatica*). Data from Wang et al. (2024).

#### Larval mortality

To investigate the positive or negative effects of selected phenolic PSMs on NPV-mediated control of *S. exigua* larvae, the mortality of larvae exposed to NPV and PSMs (genistein, kaempferol, quercetin, or coumarin) was assessed. Third-instar larvae were fed artificial diet cubes containing PSM [0.1% (186 mg/larva)] and SeMNPV [2.12 × 10^3^OBs·mL^-1^ (2.12 OBs/larva)] in a factorial design. The PSM treatments received diet cubes containing a single PSM or no PSM; the NPV treatment groups received diet cubes with 1 mL of NPV suspension [2.12 × 10^3^OBs·mL^-1^ (2.12 OBs per larva)] or sterile water as a control. Following NPV exposure (or feeding on the NPV-free control diet), larvae continued to be fed with NPV-free artificial diet containing the same type and dose of PSM as the initial treatment until pupation. Larval mortality was observed and recorded daily. The corrected mortality rate was calculated using the following formula: [(Treatment group mortality – Control group mortality) / (100 – Control group mortality)] × 100%. Data from Wang et al., (2024).

#### Hemolymph collection

Hemocytes from larvae were collected using capillaries (Wang et al., 2021). In the early stage (at 24 hours post-treatment), the larvae were smaller, and we collected 1.0-2.0 μL of hemolymph from each larva. When the larvae were larger, 48 and 72 hours post-treatment, we collected 2.0-4.0 μL of hemolymph from each larva. Specifically, one pool consisted of hemolymph from 10 to 20 larvae at 24 hours post-treatment, while another pool was from five to ten larvae at 48 and 72 hours post-treatment. Three pools, serving as biological replicates, were used for seven different analyses: morphological observation of hemocytes (4 groups × 3 time points × 3 replicates with 10 larvae per sample), apoptosis rates (4 groups × 3 time points × 3 replicates with 10 larvae per sample + 9 groups × 3 replicates × 10 larvae per sample), transcriptome sequencing (4 groups × 3 replicates with 10 larvae per sample), gene expression measurement of microRNAs (miR-8, miR-6497-3p, and miR-9c-5p) (4 groups × 3 time points × 3 replicates with 10 larvae per sample + 15 groups × 3 replicates with 10 larvae per sample), and mRNAs (MAPK, ASK1, and 14-3-3 protein zeta) (4 groups × 3 time points × 3 replicates with 10 larvae per sample + 15 groups × 3 replicates with 10 larvae per sample), enzymatic activity (MAPK, ASK1, and 14-3-3 protein zeta) (3 proteins × 4 groups × 3 time points × 3 replicates with 10 larvae per sample), and gene expression measurement of microRNAs (miR-8, miR-6497-3p, and miR-9c-5p) (4 groups × 3 time points × 3 replicates with 10 larvae per sample) and mRNAs (MAPK, ASK1, and 14-3-3 protein zeta). In total, approximately 3810 larvae (630 + 360 + 120 + 810 + 810 + 1080) were used in this experiment.

#### Configuration of hemocyte observation

Scanning electron microscopy was utilized to observe the morphology of hemocytes. A volume of 20 μL of hemolymph was collected and centrifuged at 200 × g for 2 minutes, after which the supernatant was discarded. The cells were resuspended in 500 μL of PBS and placed onto a glass culture dish. Subsequently, 4% glutaraldehyde was slowly added to the edge of the dish, which was gently swirled for 10 seconds to soak the cells. After fixing the cells for 2 hours, the glutaraldehyde was removed, and the hemocytes were washed three times with PBS, each wash lasting 15 minutes. The samples underwent dehydration using tert-butanol. They were successively immersed in glass culture dishes containing 30%, 50%, 70%, 80%, 90%, and 100% tert-butanol solutions for 5-minute intervals of gradient dehydration. After the stepwise dehydration process, the samples were immersed in 100% pure tert-butanol once more and placed in a Petri dish at 4 °C for 15 minutes. The samples were then transferred to a vacuum dryer and dried on a low setting. The dried samples were coated with gold using an ion sputtering apparatus and observed under a scanning electron microscope at 10 kV. A total of 360 larvae (divided into 4 groups with 3 replicates, each containing 10 larvae) were used for the assay.

#### Apoptosis rate measurement

At each of the three post-treatment time points, 20 μL of hemolymph was collected from each replicate and then centrifuged (200 × g at 4 °C for 5 minutes, with the supernatant discarded). Hemocytes were collected at a concentration of 1-5 × 10^5^ and resuspended in 500 μL of PBS. The cells were gently swirled, centrifuged again (200 × g at 4 °C for 5 minutes, with the supernatant discarded), and then resuspended in 500 μL of binding buffer (containing 10 mmol·L^-1^ Hepes/NaOH, pH 7.4, 140 mmol·L^-1^ NaCl, and 2.5 mmol·L^-1^ calcium chloride). Subsequently, 5 μL of Annexin v-FITC and 5 μL of propidium iodide (PI) were added to the cell suspension and mixed thoroughly. The cells were then analyzed by flow cytometry after being allowed to react at room temperature (20-25 °C) in the dark for 15 minutes. The apoptosis rate was calculated as the sum of the early and late apoptosis rates in this study. A total of 360 larvae (comprising 4 groups with 3 replicates, each containing 10 larvae) were used for the assay.

#### Transcriptome sequencing

In the study of *S. exigua*, transcriptome sequencing was employed to identify differentially expressed miRNAs in larvae treated with NPV, coumarin (C), NPV+coumarin (C), and a control (CK). Based on our previous research, an NPV treatment level of 2.12 × 10^3^ OB·mL^-1^ and a coumarin dosage of 0.01% dw were selected for the experiment (Wang et al., 2024). The same dosages of NPV and coumarin were used in subsequent tests.

Total RNA was isolated using the Trizol Reagent (Invitrogen Life Technologies), and its concentration, quality, and integrity were determined using a NanoDrop spectrophotometer (Thermo Scientific). Small RNA libraries were constructed using the NEBNext Multiplex Small RNA Library Prep Set for Illumina (New England Biolabs, Inc.) following the manufacturer’s instructions. Briefly, 1μg of total RNA from each sample was ligated to a 3’ adapter and a 5’ adapter using a Ligation Enzyme Mix. The resulting samples were reverse-transcribed using Superscript II reverse transcriptase, and PCR products were amplified. All steps were performed according to the manufacturer’s protocols. Small RNA libraries were analyzed for quality control, and the average size of inserts was approximately 140 to 150 bp. The sequencing library was quantified using the Agilent high sensitivity DNA assay on a Bioanalyzer 2100 system (Agilent) and then sequenced on the NovaSeq 6000 platform (Illumina) by Shanghai Personal Biotechnology Cp. Ltd.

The quality information of raw data in FASTQ format was calculated, and the raw data was filtered using a self-developed script by Personalbio company, resulting in clean data by removing adapter and low-quality sequences. Clean Reads ranging from 18 nt to 36 nt in length were filtered and deduplicated to obtain Unique Reads for subsequent analysis. A reference genome index was built using Bowtie2 (v2.5.1), and the de-duplicated Clean Reads were mapped to the reference genome using miRDeep2 (v2.0.0.8) software. Unique Reads were annotated with known miRNAs from the miRBase database (http://www.mirbase.org/) and then with other non-coding RNAs.

A family of known miRNAs (derived from miRBase) was classified, and the presence of the corresponding miRNA family in other species was investigated. The Reads Count value of the miRNA was counted based on the number of sequences aligned to the mature miRNA. The highest abundance of miRNA with the same name was chosen for subsequent analysis. Differential expression analysis of miRNAs was conducted using DESeq (v1.39.0), and transcripts with |log2FoldChange|>1 and P-value<0.05 were considered differentially expressed miRNAs.

The R language Pheatmap (v1.0.12) software package was used to perform bidirectional cluster analysis on all miRNAs and samples. The Euclidean method was used to calculate the distance, and the hierarchical clustering longest distance method (Complete Linkage) was used for clustering.

The 3’UTR sequence of the mRNA of this species was considered the target sequence, and MiRanda (v3.3a) was used to perform target gene prediction on the differentially expressed miRNA sequence. Gene ontology (GO, http://geneontology.org/) and Kyoto Encyclopedia of Genes and Genomes (KEGG, http://www.kegg.jp/) enrichment analysis were performed on the target genes of differentially expressed miRNA. TopGO (v2.50.0) was used to perform GO enrichment analysis on the target genes of differential miRNA, calculating the P-value by the hypergeometric distribution method (the standard of significant enrichment is *P*-value <0.05), and identifying the GO term with significantly enriched differential genes to determine the main biological functions performed by differential genes. The clusterProfiler (v4.6.0) software was used to carry out the enrichment analysis of the KEGG pathway of the target genes of differential miRNA, focusing on the significant enrichment pathway with P-value <0.05. In total, 120 larvae (4 groups × 3 replicates with 10 larvae per sample) were used for this test.

#### qRT-PCR analysis

Total RNA was extracted following the previously outlined protocol. Primers, detailed in Table S3, were created employing the Primer5 software and subsequently synthesized by Sangon Biotech (Shanghai) Co., Ltd., China. For the quantitative real-time PCR (qRT-PCR), Green miRNA Two-Step qRT-PCR SuperMix (designed for microRNA genes) and TransStart Top Green qPCR SuperMix (intended for mRNAs) were utilized with a Step-One System from Trans Gen Biotech, located in Haidian District, China. The qRT-PCR procedure included an initial denaturation phase at 94°C lasting 5 minutes, succeeded by 35 cycles comprising denaturation at 94°C for 30 seconds, annealing at 55°C for 30 seconds, and extension at 72°C for 2 minutes. A final extension at 72°C for 10 minutes was performed to assess the expression of the MAPK, ASK1, and 14-3-3 protein zeta genes. Expression levels of MAPK, ASK1, and 14-3-3 protein zeta were quantified utilizing a qRT-PCR protocol characterized by the following cycling conditions: initial denaturation at 94°C for 30 seconds, followed by 45 cycles of denaturation (94°C for 5 seconds) coupled with annealing (60°C for 30 seconds), and a final extension phase (72°C for 30 seconds).^12,63^ Gene expression analysis was carried out using the 2-ΔΔCt method as described by Livak and Schmittgen.^64^ The β-actin or U6 gene from *S. exigua* served as the reference gene, with the expression level of uninfected *S. exigua* larvae fed an artificial diet established as the standardized value of 1. The 2-ΔΔCt method was employed to evaluate the expression levels of the target genes. In total, 720 larvae (2 test × 4 diets × 3 time points × 3 replicates with ten larvae per sample) were analyzed for gene expression.

#### Assay of enzyme protein levels

The evaluation and quantification of mitogen-activated protein kinase (MAPK), apoptosis signal-regulating kinase 1 (ASK1), and 14-3-3 protein zeta protein levels were conducted following the guidelines provided by the Enzyme-Linked Immunosorbent Assay kit (Shanghai Fantai Biotechnology Co., Ltd., Shanghai, China), using the specific techniques outlined by Wang et al.^12^ An ANOVA was also employed to ascertain if the protein levels exhibited significant dependence (*P* < 0.05) on the predetermined variables: larval diet (coumarin or coumarin-free) and viral treatment (NPV-infected or uninfected). In total, 1080 larvae (3 proteins × 4 diets × 3 time points × 3 replicates, with ten larvae per sample) were utilized in this experiment.

#### Dual luciferase reporter assay

To further validate the prediction results of the miRNAs, a dual-luciferase assay was conducted to verify the regulatory interaction between miRNAs (miR-8, miR-6497-3p, and miR-9c-5p) and their potential target genes (MAPK, ASK1, and 14-3-3 protein zeta). The sequences of MAPK, ASK1, 14-3-3 protein zeta, as well as those of miR-8, miR-6497-3p, and miR-9c-5p, along with their respective mutant variants lacking docking sites, were synthesized. These sequences were then sub-cloned into luciferase reporter vectors, designated as MAPK-wt, MAPK-mu, ASK1-wt, ASK1-mu, 14-3-3 protein zeta-wt, 14-3-3 protein zeta-mu, miR-8-wt, miR-8-mu, miR-6497-3p-wt, miR-6497-3p-mu, miR-9c-5p-wt, and miR-9c-5p-mu. Subsequently, sequencing was performed to confirm these plasmids. Additionally, the relative activity of the luciferase enzyme was measured using the Dual-Luciferase Assay Kit (Wuhan GeneCreate Biological Engineering Co., Ltd., China), strictly following the manufacturer’s protocol.

#### Identification of miRNA and their target genes in vivo

Agomirs or antagomirs of microRNAs (miR-8, miR-6497-3p, and miR-9c-5p) were designed and synthesized by Sangon Biotech (Shanghai) Co., Ltd., China (Table S4). Following treatment of the larvae with NPV for 48 hours, one microliter of agomir, antagomir, or siRNA (0.5 μg·μL^−1^) was injected into the abdomen of early third-instar larvae between the third and fourth abdominal segments using a microsyringe (Zinsser Analytic). After injection, active larvae were selected for RT-PCR analysis. Each sample was ground to a fine powder in liquid nitrogen and then centrifuged at 1000 × g for 20 minutes. Subsequently, the expression of miRNAs (miR-8, miR-6497-3p, miR-9c-5p) and mRNAs (MAPK, ASK1, 14-3-3 protein zeta) was determined by RT-PCR analysis as described above [N = 270 larvae, (9 groups × 3 replicates × 10 larvae].

#### RNA interference and examination of the gene function

Small interfering RNAs (siRNA) were also designed and synthesized by Sangon Biotech (Shanghai) Co., Ltd., China (Table S6). Following treatment of the larvae with NPV for 48 hours, one microliter of siRNA (1μg·μL^−1^) of interest or negative control (NC) siRNA (Sangon Biotech) was injected. Post-injection, active larvae were selected for subsequent experiments. Gene expression was determined using Real-Time PCR analysis, as previously described. The gene expression level of the NC-injected larvae at 4 hours was set as the baseline (1), and the expression levels of other genes were compared accordingly. Samples were collected at 4, 12, and 24 hours post-injection (N = 270, calculated as 5 larvae × 3 genes × 3 time points × 2 siRNA treatments × 3 replicates). Subsequently, the apoptosis rate (N = 960, calculated as (16 groups × 3 replicates × 10 larvae) and the mortality rate of the larvae were measured at selected time points post-injection (N = 1920, calculated as (16 groups × 3 replicates × 40 larvae).

## QUANTIFICATION AND STATISTICAL ANALYSIS

The normal distribution and homoscedasticity of all data were assessed using the Kolmogorov-Smirnov test and the Levene tests. A three-way analysis of variance with a General Linear Model was employed to analyze the effects of three factors (NPV, coumarin, and time after treatment) on apoptosis rates, the expression of three microRNA genes (miR-8, miR-6497-3p, and miR-9c-5p), and the activities and mRNA expressions of three proteins in *S. exigua* larvae (MAPK, ASK1, and 14-3-3 protein zeta). Statistical analyses in this study were conducted using SAS 9.4 (SAS Viya). Pearson correlations were used to analyze the relationships among variables. Data are presented as means ± SEM. Differences among groups were calculated using ANOVA, and Tukey’s HSD post hoc tests were applied to determine significant mean differences (*P* < 0.05).

